# REVA as a Well-curated Database for Human Expression-modulating Variants

**DOI:** 10.1101/2021.02.24.432622

**Authors:** Yu Wang, Fang-Yuan Shi, Yu Liang, Ge Gao

## Abstract

More than 80% of disease- and trait-associated human variants are noncoding. By systematically screening multiple large-scale studies, we compiled REVA, a manually curated database for over 11.8 million experimentally tested noncoding variants with expression-modulating potentials. We provided 2424 functional annotations that could be used to pinpoint plausible regulatory mechanism of these variants. We further benchmarked multiple state-of-the-art computational tools and found their limited sensitivity remains a serious challenge for effective large-scale analysis. REVA provides high-qualify experimentally tested expression-modulating variants with extensive functional annotations, which will be useful for users in the noncoding variants community. REVA is available at http://reva.gao-lab.org.

## Introduction

Noncoding regions occupy the majority of the human genome [1]. It has been demonstrated that noncoding variants can affect the regulation of genes [2] and more than 80% of disease- and trait-associated variants are noncoding variants [3]. Noncoding variants that could affect gene expression can be considered as expression-modulating variants [4]. Several experimental assays have been developed to characterize expression-modulating variants. Genome editing technologies such as transcription activator-like effector nucleases (TALENs), zinc finger nucleases (ZFNs), and clustered regularly interspaced short palindromic repeats with Cas9 nuclease (CRISPR/Cas9) provide high-quality validated data but are generally low throughput [5–7]. Recently developed massively parallel reporter assays (MPRAs) can identify transcriptional regulatory elements in an efficient way, allowing systematic screening of tens of thousands of genetic variants for pinpointing the causal variants of complex traits [4, 8, 9]. All expression-modulating variants stored in MaveDB [10] are validated by the MPRA experiments. MPRAs have generated over 10 million human expression-modulating variants [11], however, only around 30 thousand of them have been collected by MaveDB without any functional annotation, which hinders the further utilization of these data.

Though experimental assays for characterizing noncoding expression-modulating variants have generated huge amount of data, it is still inadequate for covering all noncoding variants identified in human genomes. Therefore, multiple computational tools have been developed for identifying expression-modulating variants (**Table 1**). Transcription factors (TFs) could regulate genes through binding to sequence motifs [12] and noncoding variants could affect gene regulation by changing motifs [13]. Funseq2 integrated a module for detecting motif-breaking and -gain events through the change of position weight matrix (PWM) and other functional annotations to prioritizing cancer driver mutations [14]. Methods based on machine learning have been used wildly in biological researches [15]. CADD [16] used support vector machine (SVM) to classify variants into functional and nonfunctional variants and GWAVA [17] used random forest to predict disease-related variants. Both of CADD and GVAWA are based on supervised learning methods, while Eigen [18] implemented unsupervised learning methods to classify variants. All these tools highly depend on existing annotations at corresponding loci. In 2015, Alipanahi et al. [19] developed DeepBind based on convolutional neural networks (CNNs) to predict the binding affinity between TFs and DNA or RNA binding proteins and RNA. DeepSEA [20] applied similar methods to predict the effect of noncoding variants on binding affinity and then classify variants through logistic regression into functional or nonfunctional groups. All tools mentioned above identifying expression-modulating variants are through indirect inference, because they are not trained on expression-modulating variants or expression-related data. EnsembleExpr [21] used MPAR data for training an ensemble-based model to characterize expression-modulating variants directly. ExPecto [22] *ab initio* predicts the variants’ effects on gene expression from 40kb promoter-proximal sequences and then pinpoints expression-modulating variants. However, there is no comprehensive evaluation of these computational tools based on high-quality expression-modulating variants, therefore, it is difficult for users to choose appropriate tools for their tasks.

**Table 1.**
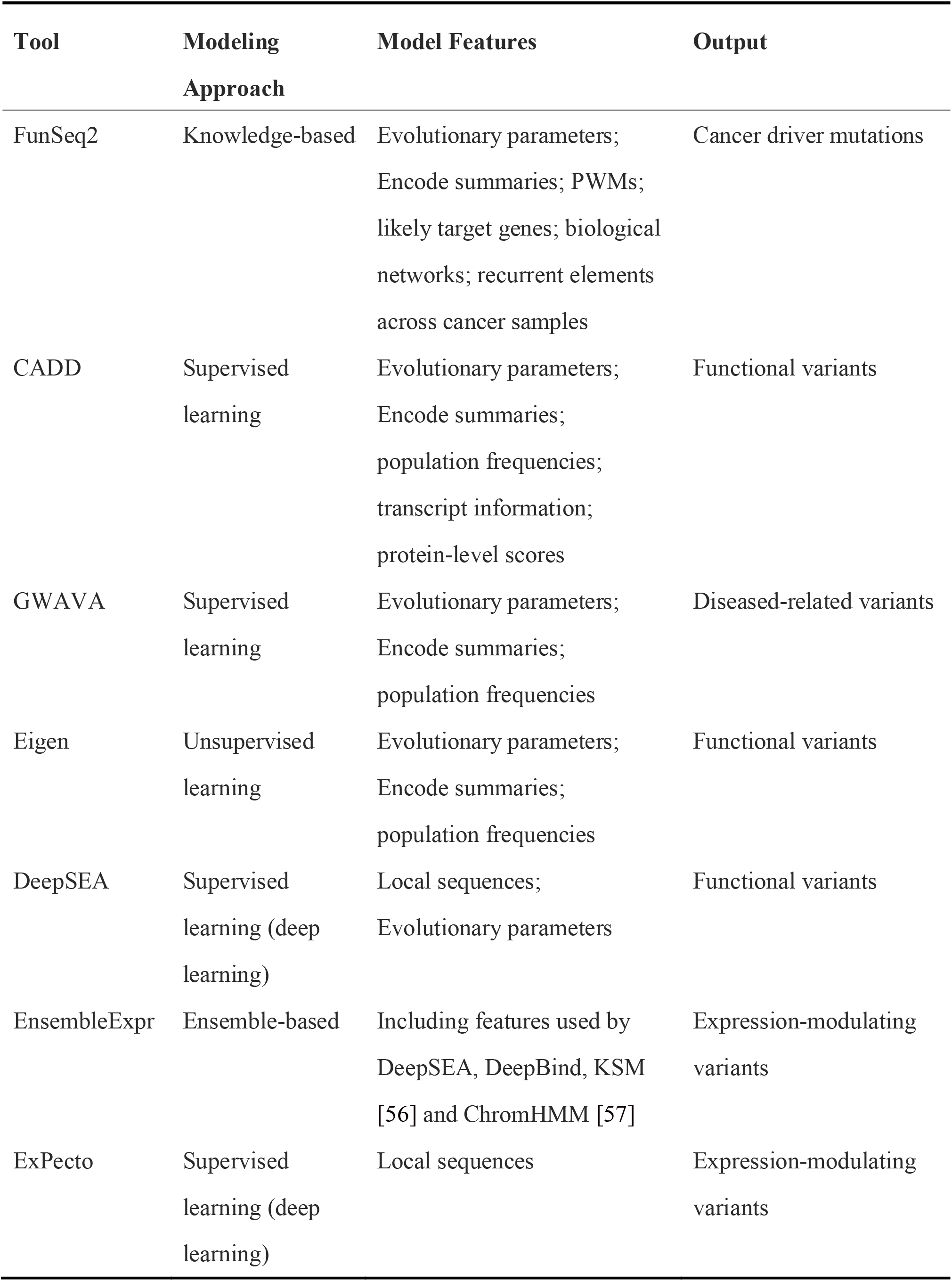
Properties of involved computational tools.

Here, we present a repository for <u>e</u>xpression-modulating <u>va</u>riants (REVA). The current release of REVA consists of over 11.8 million experimentally validated expression-modulating variants in the human genome, curated with extensive functional annotations. We further benchmarked seven popular computational tools in identifying expression-modulating variants [14, 16–18, 20–22] based on high-quality data in REVA. All data and benchmarking results are publicly available at http://reva.gao-lab.org.

### Construction and content

#### Data collection and integration

To ensure unified and high-quality data, all records in REVA were collected and curated using a standard procedure (**Figure 1**). We used a list of keywords, “MRPA”, “STARR-seq”, “CRE-seq” with “mutation”, “variant” and “variation”, to obtain publications from PubMed (https://pubmed.ncbi.nlm.nih.gov/) and then manually checked the abstracts and full texts of the matching publications to obtain literatures that experimentally validated the effects of noncoding expression-modulating variants.

**Figure 1.**
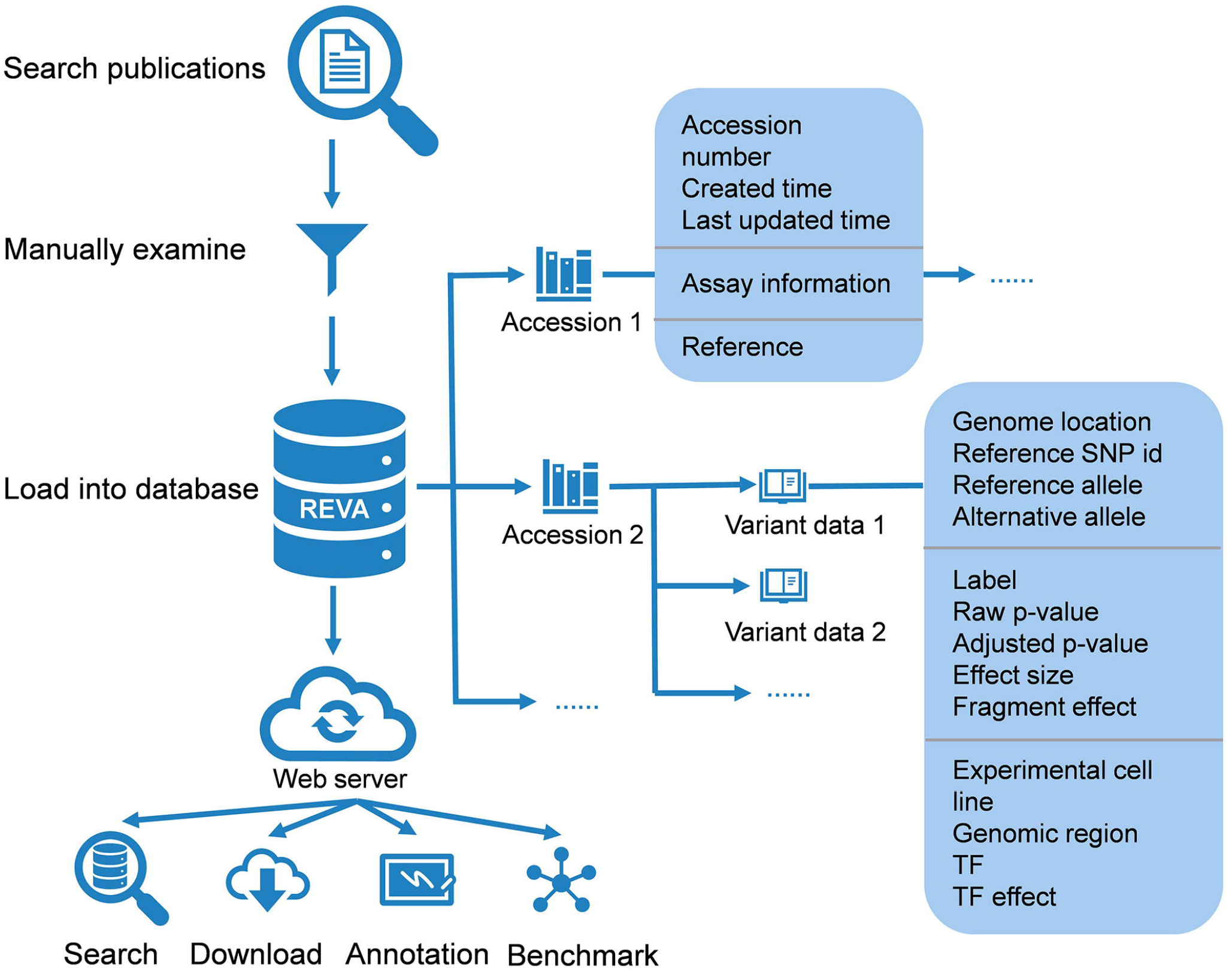
Overview of the structure of REVA. Manually curated noncoding variant data as well as supplemental information were stored in the database at two levels: accession and variant data. Accession contained the information about the publication, and variant data contained all related information about the variant. A web interface was built for users to access the data in the database.

For filtered literatures, we extracted related information of the variants from the main text as well as supplementary materials of publications and converted them to the same format (**Table 2**). Variants that failed to be mapped to both GRCh37 and GRCh38 were removed. Variants only mapped to coding region were also removed. For missing information, we used “.” as a placeholder. In addition, the detail protocol and raw data of the experiments were also extracted.

**Table 2.**
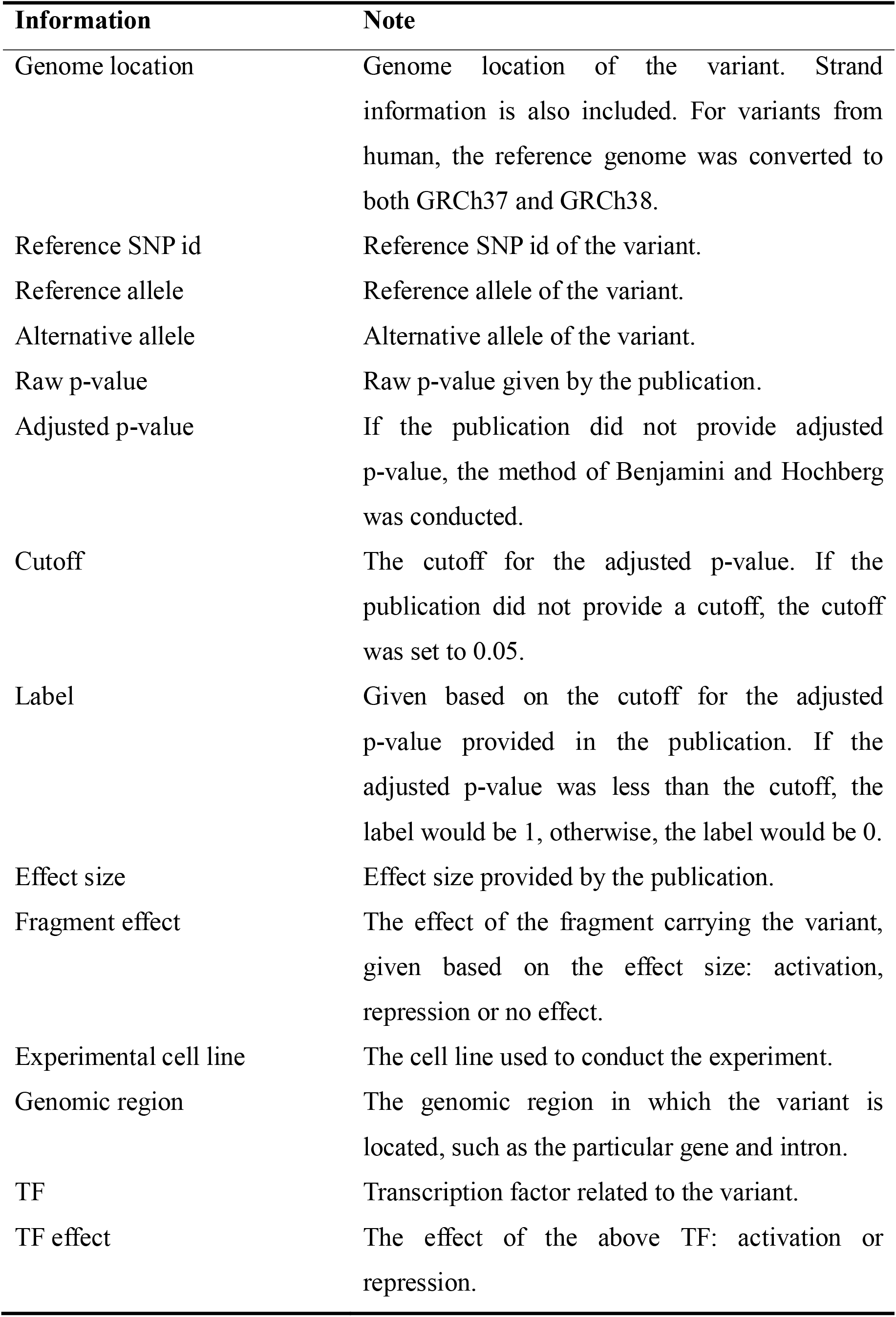
Variant information extracted during the data collection process.

For variant data with the same chromosome, genome location, reference allele, alternative allele and experimental cell line from different publications, a meta-analysis was conducted to integrate data. The harmonic mean p-value (HMP) method [23] was used in the meta-analysis, and the cutoff for the meta p-value was set to 0.001 to generate the meta-label. The variants involved in the meta-analysis but without a raw p-value were also removed.

The label of variants was given based on the cutoff for the adjusted p-value or meta p-value, and then variants were classified into positive variants and negative variants based on label or meta-label. If the variant’s label was 1, the variant was a positive variant and considered to have effects on gene expression, otherwise, it was a negative variant without effect on gene expression.

#### Database construction

All manually curated variant data as well as meta-information were stored in MongoDB (https://www.mongodb.com/) at two levels: accession and variant data (Supplementary Figure S1). Each accession entry consisted of an accession number, the times of creation and last update for the accession, information about the assay used in the publication (method type, original reference genome version, link to raw data, and summary of the assay), and the reference. Variant data include all related information about the variant, and each variant data entry is linked to one accession. For the data involved in the meta-analysis, the variant data contained the results of the meta-analysis and linked to all related variants and accessions.

We also integrated DisGeNET v7.0 (https://www.disgenet.org/) [24] variant-disease associations, GWAS catalog (https://www.ebi.ac.uk/gwas/), ClinVar (https://www.ncbi.nlm.nih.gov/clinvar/), COSMIC (https://cancer.sanger.ac.uk/cosmic), and three-dimensional interacting genes and chromatin state from 3DSNP [25] to our database for providing more variant information.

#### Variant annotation

In efforts to pinpoint plausible regulatory mechanisms for these variants, we used 2403 trained CNNs to annotate the functional effects of sequence variations [26] based on 1249 TF binding, 766 histone modification, 280 DNA accessibility, and 108 DNA methylation profiles from recent Encyclopedia of DNA Elements (ENCODE) data.

An NVIDIA Tesla P100 Graphics Processing Unit with the implementation on the deep learning framework TensorFlow (https://www.tensorflow.org/) and Python (https://www.python.org/) were used for training models. We adopted stochastic gradient descent (SGD) as the optimizer and the initial learning rate was 0.01.

The final output layer of CNN model was a fully connected layer with sigmoid function used to scale the output between 0 to 1. The input layer was one dimensional convolution layer with threshold ReLU as the activation function. Next, the max pooling layer was performed to reduce the complexity of data. Then dropout layer was used to avoid overfitting problem. The next two layers were full connected layer with threshold ReLU as the activation function and dropout layer.

For TF binding, histone modification, and DNA accessibility models, the positive data for training CNNs were the 200bp sequences centered on the peak in ENCODE profiles. Then we remove positive sequences from human reference genome and split the rest to 200bp bins. Random sampled 200bp bins with the same number of positive data were used as negative data. For DNA methylation, the 200bp sequences centered on the target base with the methylation rate more than 0.5 or less than 0.5 in whole-genome bisulfite sequencing (WGBS) data were considered as positive data and negative data respectively. One-hot encoding was conducted to transformed each sequence to 200×4 binary matrix for model training.

Five-fold cross-validation strategy was used to train models. During each iteration of model training, 15% of the input data were random selected as the independent testing dataset to evaluate model performance. The remained data were split with 70% to train models and 15% as the validation dataset to optimize parameters. Model performance was evaluated with area under the receiver operating characteristic curve (AUROC) and area under the precision recall curve (AUPRC) to test the sensitivity and specificity, and models with best performance were selected for variant annotation. An average AUROC and AUPRC of 2403 models were reported.

To character the binding affinity changes of the variant, we used 2403 trained CNNs to predict on 200bp sequences centered with the reference allele and alternative allele respectively. For each chromatin profiles, the log2-fold-changes (as the method shown in DeepSEA) [20] was calculated as the variant effect on chromatin profiles. Specifically,

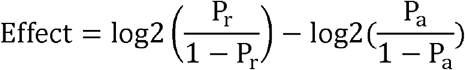

where P_r_ is the prediction of sequence with reference allele and P_a_ is the prediction of sequence with alternative allele.

Furthermore, we incorporated 13 DNA physicochemical properties and 8 evolutionary features into the annotation pipeline. The 13 physicochemical properties were calculated as the Rong Li et al. [27] described and 8 conservation scores were downloaded from UCSC Genome Browser (http://genome.ucsc.edu/).

#### Benchmarking

To prepare the benchmarking dataset for evaluating the performance of state-of-the-art computational tools in calling expression-modulating variants based on the curated data in REVA, we first excluded loci tested in mice (15,152). There are several overlapped variants between the training sets of state-of-the-art tools and the REVA benchmark dataset. If the benchmark dataset contains these variants, the performance of related tools will be overestimated. To avoid the influence of these variants and make a fair comparison, we further removed variants (47,518) that were either found in the GWAVA [17] and EnsembleExpr [21] training datasets or used to compute the empirical background distributions by DeepSEA [20]. For the remaining 5,809,991 loci (37,816 positive and 5,772,175 negative), we ran CADD (v1.4, https://cadd.gs.washington.edu/download) [16], DeepSEA (http://deepsea.princeton.edu/), EnsembleExpr (https://github.com/gifford-lab/EnsembleExpr/), and ExPecto (https://hb.flatironinstitute.org/expecto/) [22] and extracted from the precomputed score set of Eigen (v1.1, http://www.columbia.edu/~ii2135/download.html) [18], FunSeq2 (v2.1.6, http://funseq2.gersteinlab.org/downloads) [14], and GWAVA (https://www.sanger.ac.uk/science/tools/gwava) to obtain the corresponding predicted score for evaluation. The thresholds used in the evaluation were those recommended by the corresponding papers or official websites (Supplementary Table S1).

All variants in benchmark dataset were variants with expression-modulating potential. One of biological mechanisms that diseases-related or phenotype-related variants function is by having effects on gene expression regulation [28]. Pinpointing disease-related or phenotype-related variants is more useful for biomedical researches. Therefore, we further selected the GWAS, ClinVar, and HGMD subsets of benchmark dataset to test these tools’ power.

## Results

### Characterization and distribution of expression-modulating variants

All curated expression-modulating variants were validated by experiments and we applied standard data collection and integration procedure to ensure the high-quality data with unified format. By the end of November 2019, REVA consisted of 11,862,367 entries covering 5,948,789 experimentally tested noncoding loci across 18 cell cultures from 14 publications [4, 8, 11, 29–39]. We first excluded loci tested in mice (15,152) and with more than one alternative allele (26,326). Among the remaining 5,907,311 loci (34,700 positive and 5,872,661 negative), most were located in intergenic (positive: 49.96%, negative: 53.83%) and intronic (positive: 35.96%, negative: 39.62%) regions (**Figure 2**A, Supplementary Table S2). We found that both positive and negative variants were unevenly distributed on chromosomes, and no variants were located on the Y chromosome (Figure 2B, Supplementary Table S3). Specifically, fewer positive variants were located on chromosomes 1, 3, 5, 13, 14, 15, 21 and the X chromosome, and more positive variants were located on chromosomes 6, 8, 10, 11, 12, and 16–20. Fewer negative variants were located on chromosomes 9, 13, 14, 15, 21, 22 and the X chromosome, and more negative variants were located on chromosomes 1–8, 10, 11, 12, and 16–20. Biochemical activities were detected for 93.53% positive and 90.80% negative cases in at least one cell culture (Supplementary Figure S2, Supplementary Table S4, S5). Of note, more positive than negative variants were found in TF binding regions, highlighting the contribution of TF binding changes to expression modulation (Supplementary Figure S2).

**Figure 2.**
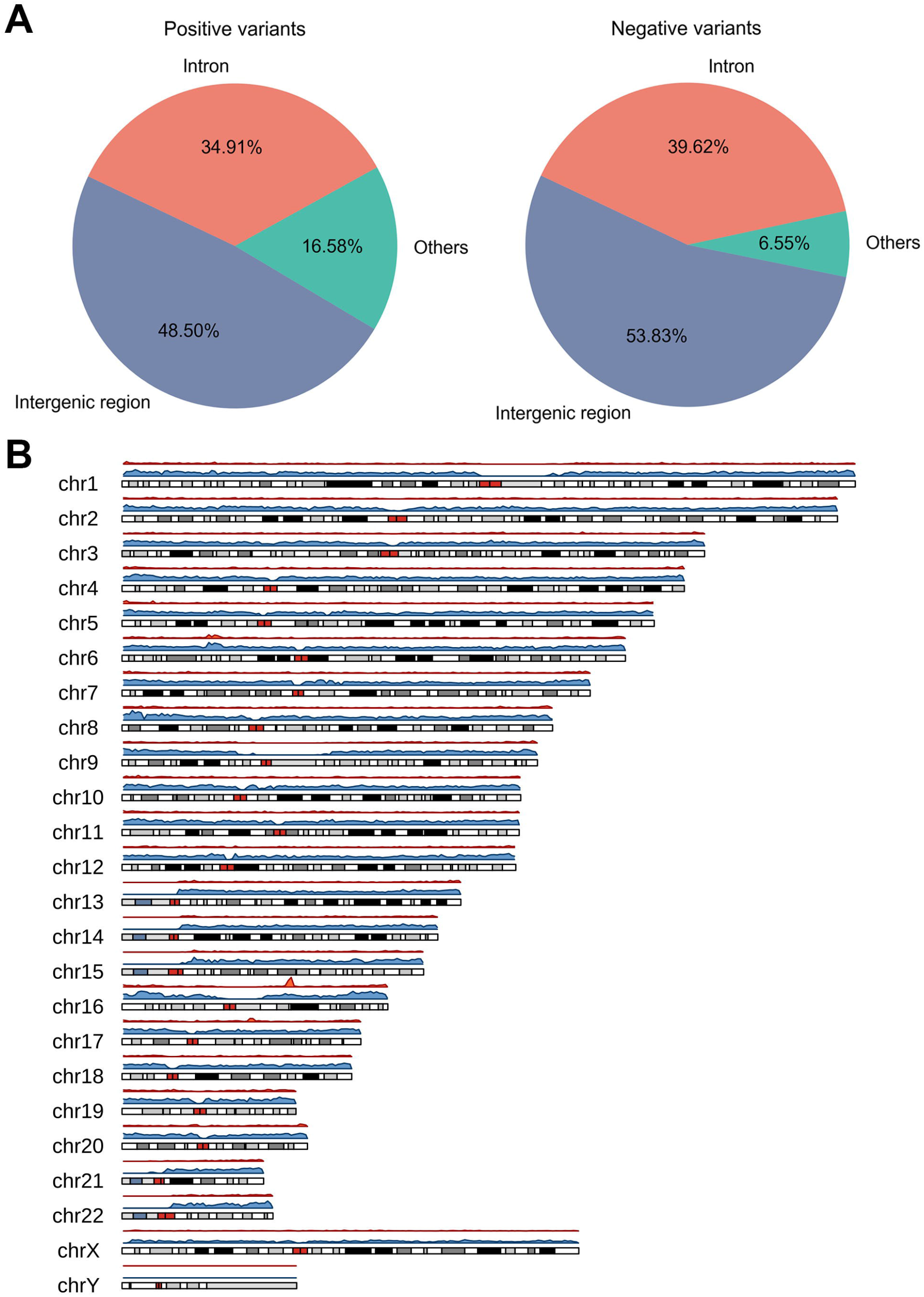
Annotation of the variants in REVA. **A**. The distribution of positive and negative variants in the genome. Mixed region refers to “with more than one type annotation”. **B**. The density distribution of positive and negative variants on chromosomes. A two-sided Fisher’s exact test with Benjamini and Hochberg correction [54] was used in the analysis of the chromosome distribution of variants. The cutoff for the adjusted p-value was set to 0.05. The density distribution plot was constructed with the karyoploteR package [55] in R. No variants were located on the Y chromosome.

### Extensive functional annotation of expression-modulating variants

We used 2403 trained CNNs to annotate the functional effects of expression-modulating variants [26]. Most of the trained CNN networks were accurate, with an average AUROC of 0.908 and an average AUPRC of 0.904. Among the 5,789,688 variants annotated, both positive and negative variants were found to lead to significant changes in binding affinity for 22 and 12 TFs on average, respectively, which also suggested that expression-modulating variants may affect gene expression regulation through changing the binding affinity of TFs. Moreover, 8.72% of positive variants and 3.56% of negative variants were located at evolutionary conserved loci (phastCons100way score > 0.6).

### Benchmarking of state-of-the-art computational tools

To evaluate the power of state-of-the-art computational tools in calling expression-modulating variants, we further benchmarked multiple state-of-the-art computational tools based on the curated data in REVA. With the benchmarking dataset containing 5,809,991 loci (37,816 positive and 5,772,175 negative), we found that 1289 could not be predicted by DeepSEA (since their evolutionary features were not available) and 560,577 were not included in the precomputed score set of Eigen, FunSeq2, and GWAVA, so we further excluded these 561,866 cases from follow-up analysis. Meanwhile, as EnsembleExpr could not finish the whole benchmarking dataset in a reasonable amount of time, we assessed its performance based on the average metrics over five randomly sampled subdatasets with 368 positive and 56,026 negative cases on average.

Overall, the best-performing tool was DeepSEA, with the highest AUROC and F1 score (**Figure 3**A and B, Supplementary Table S6). All tools performed well in terms of specificity but poorly in terms of sensitivity. EnsembleExpr had the highest sensitivity but the lowest specificity, whereas ExPecto showed the best specificity and worst sensitivity.

**Figure 3.**
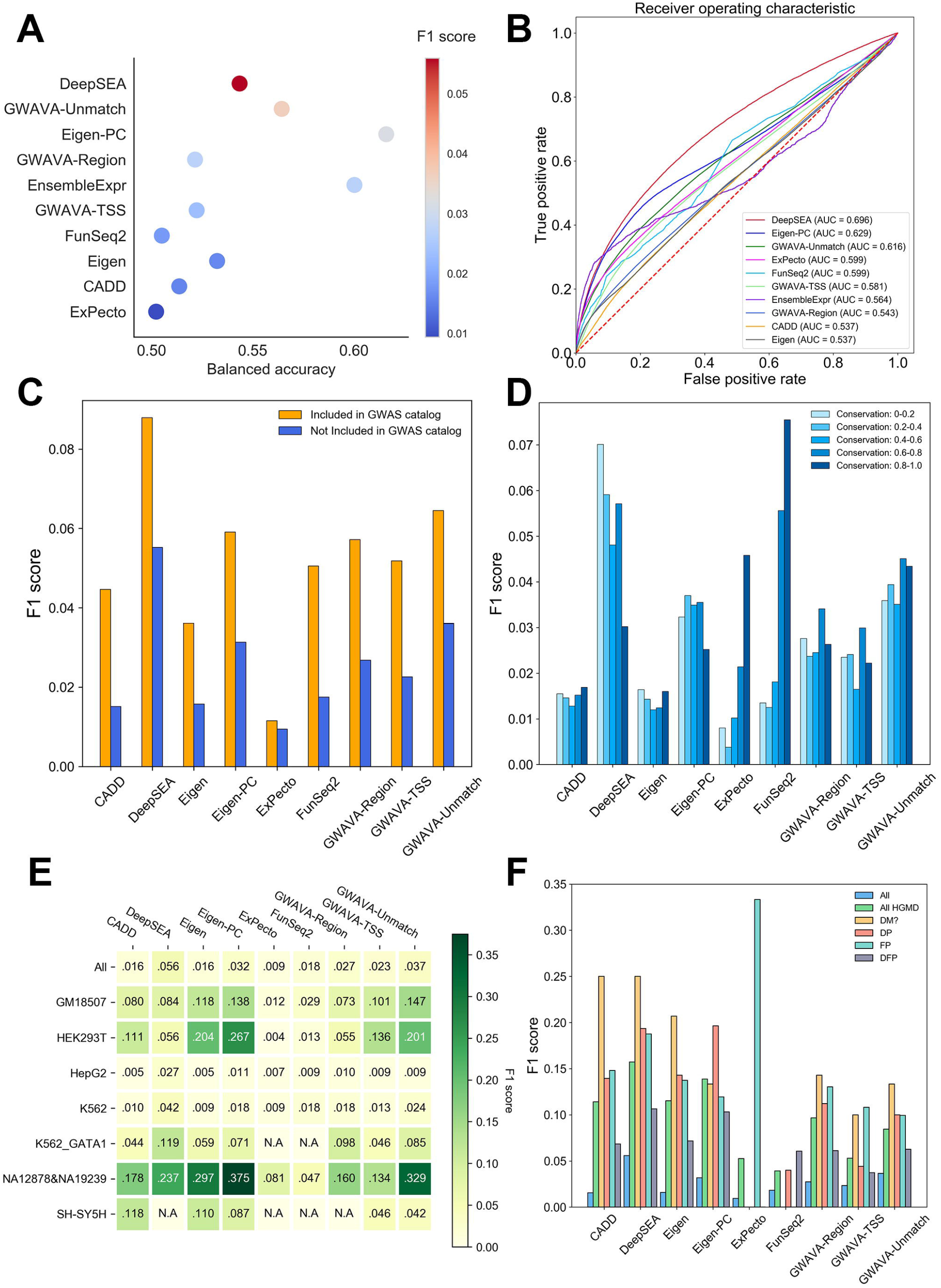
Performance of involved tools on the benchmarking dataset. **A**. The performance comparison of involved tools and bubbles colored by F1 score. The results are ordered by F1 score. **B**. The ROC curves for involved tools. **C**. Performance comparison of the involved tools except EnsembleExpr on variants that were also included in GWAS catalog and not. **D**. Performance comparison of involved tools except EnsembleExpr on variants that had different phastCons100way scores. **E**. Performance comparison of involved tools except EnsembleExpr on variants from different cell lines. “All” represents the F1 score in Figure 3A. **F**. Performance comparison of involved tools except EnsembleExpr on variants that were also included in HGMD. “All” represents the F1 score in Figure 3A. “All HGMD” represents the F1 score on all variants that were included in HGMD. “DM?”, “DP”, “FP”, and “DFP” refer to the classes of related variants documented in HGMD.

There were 52,672 variants in the benchmarking dataset that overlapped with the GWAS catalog (v1.0.2), and 658 of them were positive variants. All tools performed better on variants overlapping with the GWAS catalog, and DeepSEA still had the best performance (Figure 3C). Meanwhile, ExPecto and FunSeq2 showed better performance on variants at evolutionarily conserved loci, while DeepSEA had poorer performance (Figure 3D).

The coverage and quality of training data may contribute significantly to the performance of machine learning-based models [40]. To test whether variants from different cell lines would affect the performance of these tools, we further evaluated these tools separately on seven cell lines (Supplementary Table S7). On GM18507, GWAVA-Unmatch performed best; on HEK293T and NA12878_NA19239, Eigen-PC had the highest F1 score; DeepSEA had the best performance on HepG2, K562, and K562_GATA1; and CADD performed best on SH-SY5H (Figure 3E), which suggested that the diversity of the original training data contributed to the performance differences of these tools. Of note, thus far, only ExPecto outputs cell-type-specific scores for various tissues.

To give further explanation of the potential mechanisms of disease-related variants, we evaluated the benchmarking dataset on disease-related variants. There were 1400 variants in the benchmarking dataset that overlapped with HGMD (2019.3 professional) and 69 of them were positive variants (Supplementary Table S8). 8 of 69 variants were verified to regulated gene expression by independent experiments. 40 of 69 variants were associated with diseases such as colorectal cancer, nervous system diseases, and autoimmune diseases. To test computational tools’ power on disease-related variants, we compared their performance on these variants. All tools performed better on HGMD variants and DeepSEA still had the best performance (Figure 3F), same on variants with class “DM?” and “DFP”. Eigen-PC showed better performance on variants with class “DP”. Interestingly, ExPecto performed best on variants with class “FP” but worst on variants with other classes. We also evaluated on variants overlapped with ClinVar (2019.10.08), and DeepSEA had the best overall performance and Eigen showed better performance on “Drug response” related variants (Supplementary Figure S3, Supplementary Table S9).

### Web interface

REVA (http://reva.gao-lab.org) provides an interactive Web interface for users to explore all data entries and analysis results (**Figure 4**, Supplementary Figure S4). Users can start a quick search by chromosome position, rs id, gene name, ensembl gene id, or disease name. “Advanced search” provides a customized search and batch search for users. The query result is presented as a table, which includes basic information, expression information (such as the label, effect size and adjusted p value), and the related genomic region. Users can directly click the link of position and rs id to access UCSC Genome Browser and dbSNP (https://www.ncbi.nlm.nih.gov/snp/) for more information. Users can also click the “details” link for more information. The detail page contains eight modules: “Basic information”, “Cell Line and Expression”, “Three-dimensional Interacting Gene”, “Chromatin State”, “Disease and Phenotype”, “Meta Sources” (only available for variants involved in meta-analysis), “Accession”, and “Annotation”. In the “Annotation” module, chromatin profile features are rendered as a heatmap by cell line and a boxplot by category, and DNA physicochemical properties and evolutionary features are presented as responsive tables. Users can download the annotation for further analysis. Moreover, we also provide benchmarking results of state-of-the-art computational tools. Users can download all variants in REVA and the benchmarking dataset through the “Download” page.

**Figure 4.**
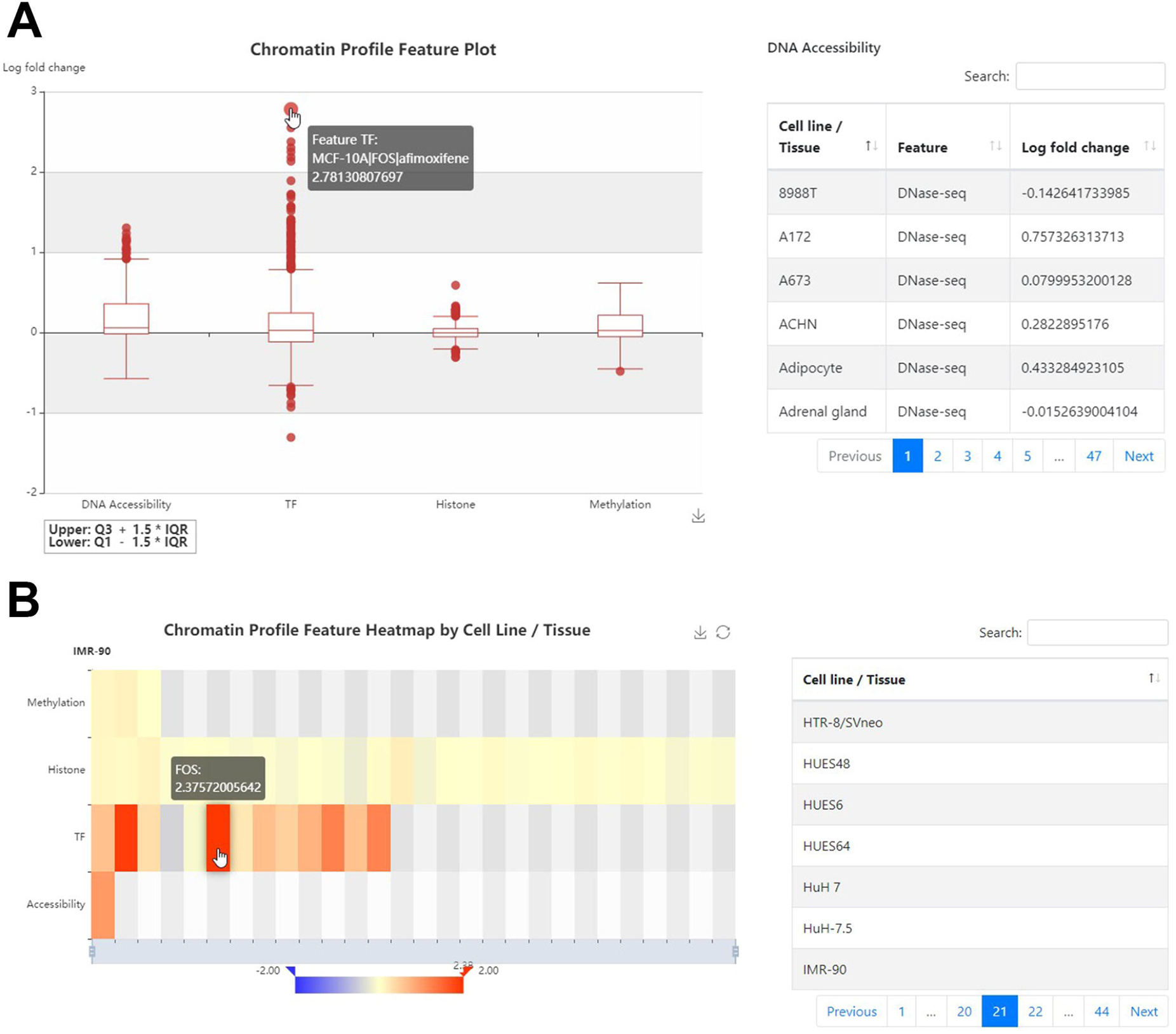
Illustration of the Web interface of REVA. **A**. Chromatin profile feature plot in “Annotation” module of variant detail page. Chromatin features are presented by category. Users can hover the mouse over the outlier or the box to show more information. At the right of boxplot is a table to show detail information. Users can click the boxplot to show corresponding category. **B**. Chromatin profile feature heatmap in “Annotation” module of variant detail page. The heatmap is presented by cell line and each row in heatmap corresponding to one category. Users can click the cell line/tissue list at the right of heatmap to render annotation in target cell line/tissue and hover the mouse over the block in heatmap to show feature information. Both Figure 4A and 4B are retrieved from http://reva.gao-lab.org/detail.php?id=intid1_8498680_8438620atk562&reference=GRCh38.

### Explore plausible regulatory mechanisms of expression-modulating variants

Autoimmune diseases are caused by abnormal immune response to attack and damage functional tissues due to complex interactions between environmental and genetic factors [41]. GWAS and fine mapping studies have identified thousands of noncoding variants associated with autoimmune diseases [42]. Since the mechanism of autoimmune disease is complicated, pinpointing causal variants and exploring their possible functional mechanism remain a challenge [43].

Ankylosing spondylitis is a kind of chronic autoimmune disease but the pathogenesis remains unclear [44]. On the advanced search page of REVA (Supplementary Figure S5), we filtered label to positive and search with “ankylosing spondylitis”. The search result contained 8 entries and among the result, the variant rs4456788 (near *ICOSLG* locus) had the largest effect size tested in HepG2 cell line and considered to repress expression. It was also tested in K562 cell line and resulting in same conclusion. Through the annotation module of the detail page, we found that in HepG2 cell line, the alternative allele of rs4456788 could decrease the binding affinity of transcription factor MAZ and FUS. MAZ has been proven to have bidirectional transcriptional regulation [45] and FUS has transcriptional activation function [46]. It could be the possible regulatory mechanism of rs445678, and this might be helpful for further researches on the mechanism of pathogenesis of ankylosing spondylitis.

## Discussion

REVA is a database specifically designed for storing experimentally validated expression-modulating data. It currently consists of 11,862,367 entries covering 5,948,789 experimentally tested noncoding loci across 18 cell cultures. Both experimentally validated expression-modulating variants and meta-information about assays were curated. Comparing with the existing database, REVA is the largest database designed for curating experimentally validated expression-modulating noncoding variants specially. Besides, we provide 2424 functional annotations including Transcription Factors binding, epigenetic modifications, DNA accessibility, DNA physicochemical properties, and evolutionary features.

Most of variants in REVA were located in intergenic and intronic regions and were unevenly distributed on chromosomes. Several factors may contribute to the uneven distributions. First, it has been well demonstrated that the functional elements are uneven distributed across chromosomes [47, 48]. Consistently, we found that the number of both positive and negative variants are highly correlated with the gene numbers across all chromosomes (Pearson’s *r* = 0.80, *p* = 2.6×10^−6^ for positive variants; Pearson’s *r* = 0.82, *p* = 7.1×10^−7^ for negative variants). Moreover, technical challenges count too. In particular, the Y chromosome had long been taken as a “genetic wasteland” [49] and excluded from genomic analyses for quite some time, due to its genetic and structure complexity [50]. Though this idea has been shifted with more researches on chromosome Y, the underrepresentation of chromosome Y on commonly used arrays still exist [51]. We also notice that certain experimental designs may also lead to reporting bias [8, 31, 36–39]. However, after removing data generated from studies designed for assessing particular regions [36] or elements [8, 31, 37–39], we found that uneven distribution remains.

Furthermore, we provide a high-quality benchmarking dataset for evaluating state-of-the-art computational tools designed for identifying expression-modulating variants as well as benchmarking results of multiple published computational tools, as a reference for users to select the best tools for their particular tasks. Overall, all seven tools have high specificity but low sensitivity. DeepSEA has the best performance on whole benchmark dataset in terms of AUROC and F1-score and all tools have better performance on disease-related or phenotype-related variants, suggesting that the diversity of the original training data of these tools contributes to different performance across different benchmark subsets. We noticed that not all tools involved in benchmark were designed for identifying expression-modulating variants originally, and a “negative” expression-modulating noncoding variant might also be associated with disease via non-transcription mechanisms like epigenetic marks [52] or chromatin structuration [53].

It should be noted that not all variants collected in our database were tested by identical experimental protocols. Non-saturation mutagenesis-based studies examine several elements at a time, and each fragment usually contains one variant, with the effect size calculated by counting reads directly [8] or employing a linear model [30]. Meanwhile, saturation mutagenesis-based studies focus on a few elements; each fragment contains two or more variants, and the effect size is calculated through linear regression [39]. Protocol details for each variant were documented during curation to help users interpret records effectively (Supplementary Figure S6).

We believe that this database will be useful for not only computational but also bench biologists in genomics, bioinformatics and genetics community, and we will keep the resources updated with new data and annotations emerged in the coming years.

## Data availability

REVA is freely accessible at http://reva.gao-lab.org.

## Code availability

Source code for all analysis and benchmark is available on GitHub at https://github.com/gao-lab/REVA-Data_Source_Code.

## CRediT author statement

**Yu Wang:** Methodology, Software, Data Curation, Formal analysis, Visualization, Writing - Original Draft, Writing - Review & Editing. **Fang-Yuan Shi:** Methodology, Software, Data Curation, Formal analysis. **Yu Liang:** Data Curation. **Ge Gao:** Conceptualization, Project administration, Supervision, Funding acquisition, Resources, Writing - Review & Editing.

## Acknowledgments

This work was supported by funds from the National Key Research and Development Program of China (2016YFC0901603), the National High Technology Research and Development Program of China (2015AA020108), as well as the State Key Laboratory of Protein and Plant Gene Research and the Beijing Advanced Innovation Center for Genomics (ICG) at Peking University. The research of Ge Gao was supported in part by the National Program for Support of Top-notch Young Professionals.

Part of the analysis was performed on the Computing Platform of the Center for Life Sciences of Peking University and was supported by the High-performance Computing Platform of Peking University.

## Supplementary material

**Figure S1.**
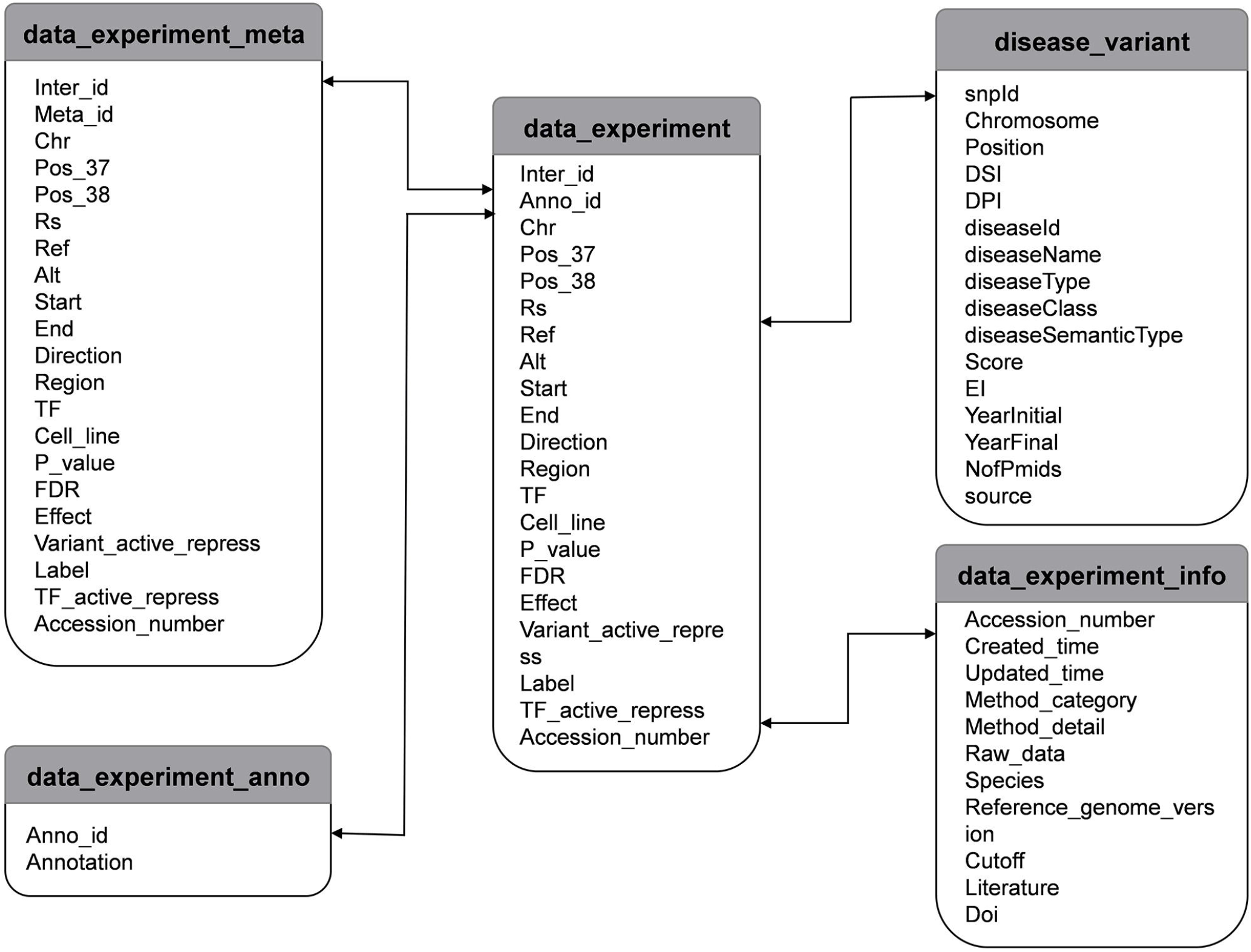
Database scheme of REVA. All data were stored in MongoDB with five collections. “data_experiment_info” contained accession information and “data_experiment” contained variant data. The original variant data included in meta-analysis stored in “data_experiment_meta”. “data_experiment_anno” and “disease_variant” contained variant annotation and associated diseases and phenotypes.

**Figure S2.**
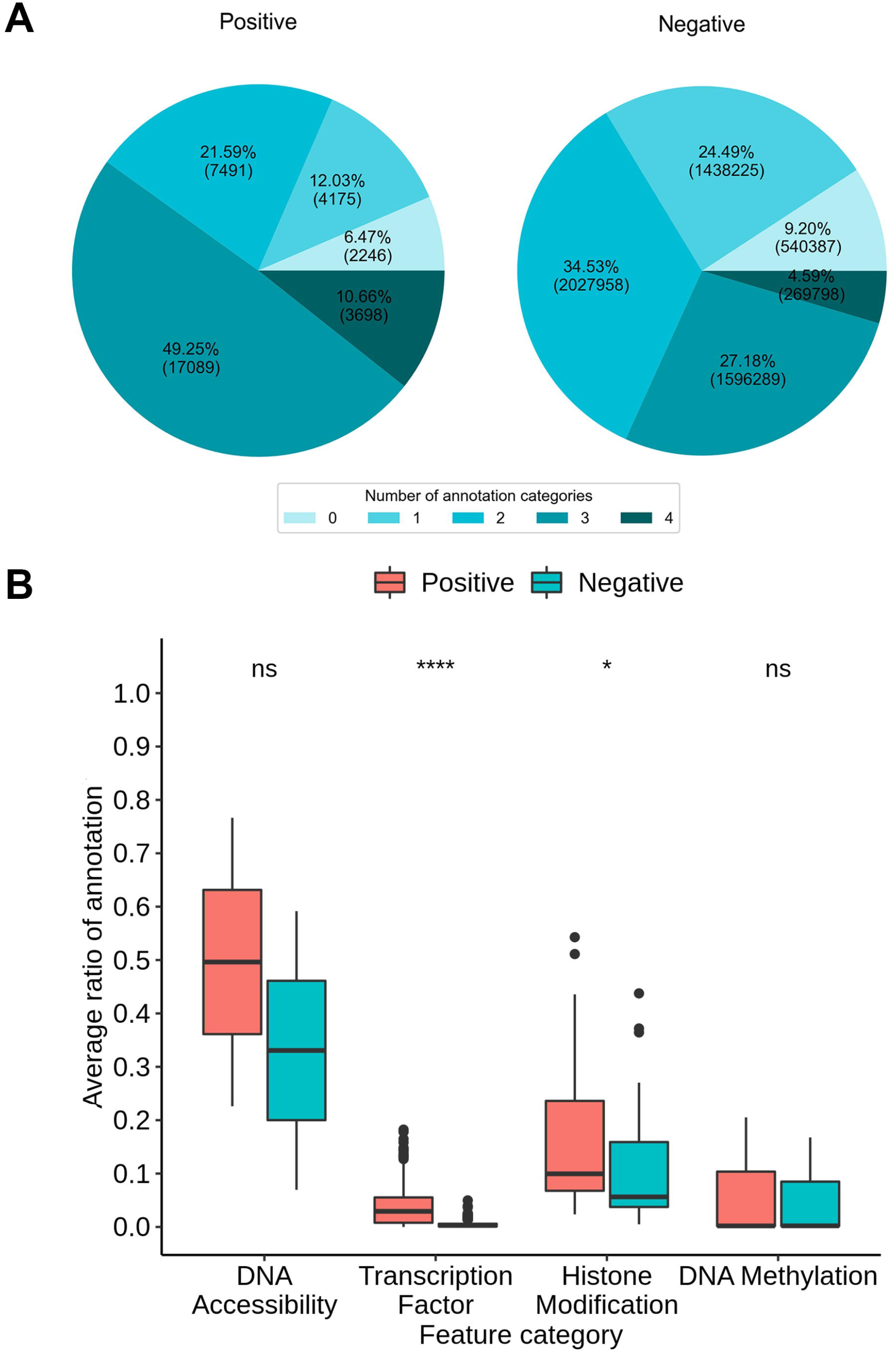
Annotation of the variants in REVA. **A**. The ratio and number of positive and negative variants that have 0, 1, 2, 3 or 4 categories of annotations. **B**. Average ratio of annotations of variants by category.

**Figure S3.**
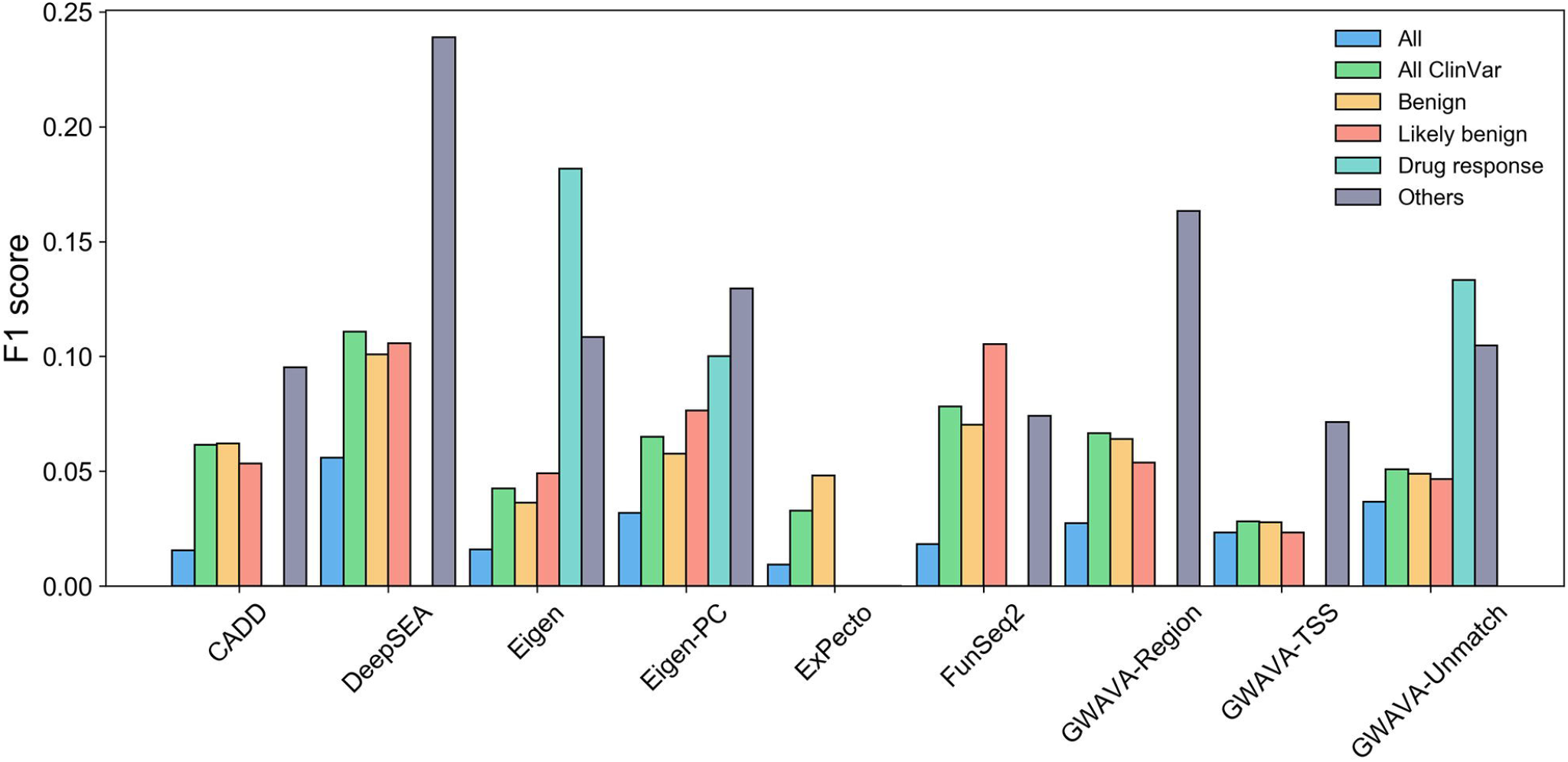
Performance comparison of involved tools except EnsembleExpr on variants that were also included in ClinVar. “All” represents the F1 score in Figure 3A. “All ClinVar” represents the F1 score on all variants that were included in ClinVar. “Benign”, “Likely benign”, and “Drug response” refer to the clinical significance of related variants documented in ClinVar.

**Figure S4.**
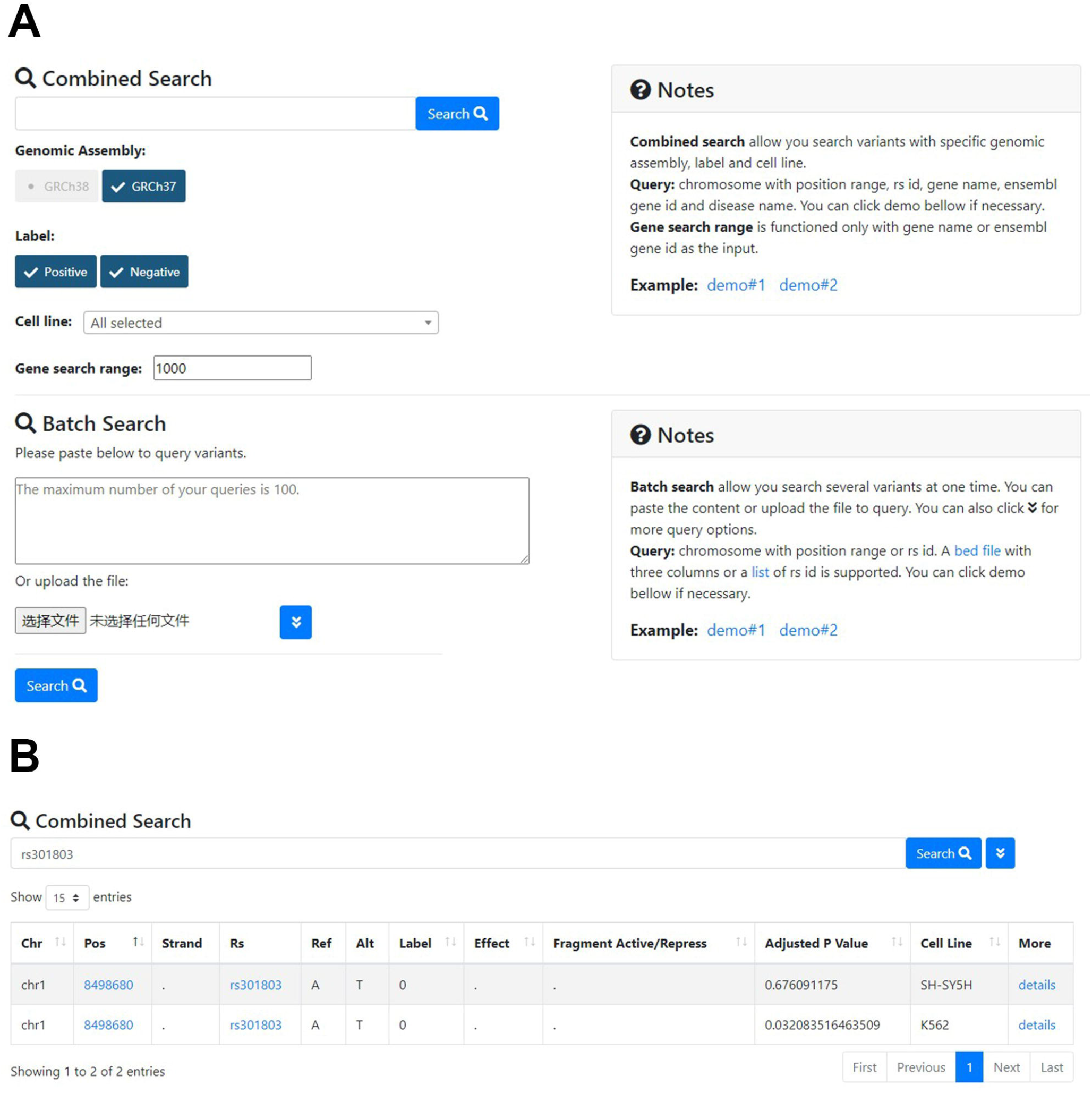
Illustration of the web interface of REVA. **A**. Advanced search page (http://reva.gao-lab.org/advanced_search.php). Users can conduct customized search or batch search through this page. **B**. Search results page. Users can sort searching results by “Chr”, “Pos”, “Label”, “Effect”, “Fragment Active/Repress”, “Adjusted P Value”, or “Cell Line”.

**Figure S5.**
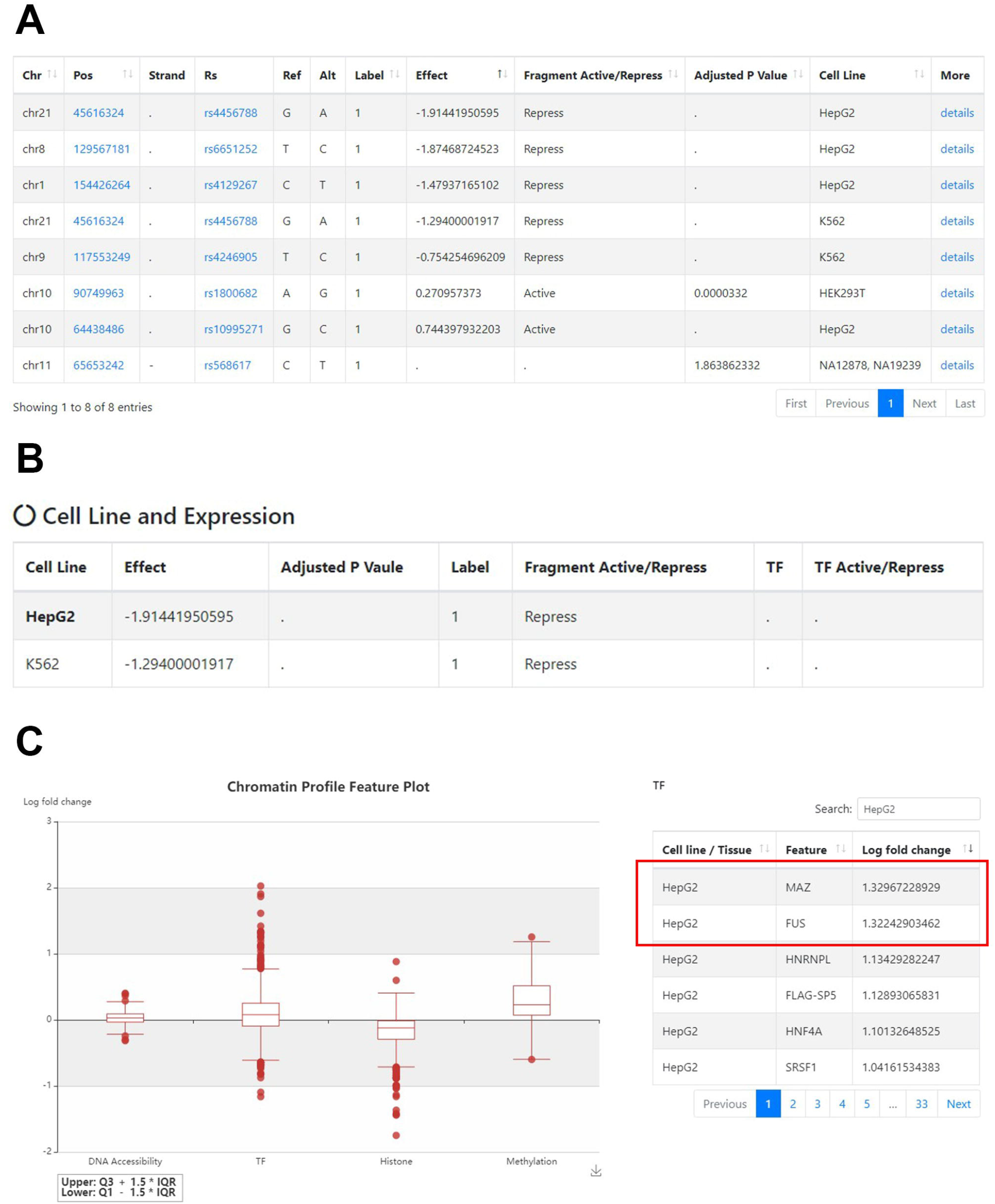
The combined search result of “ankylosing spondylitis”. **A**. Searching result sorted by “Effect”. Variant rs4456788 has the largest effect size in HepG2 cell line. **B**. “Cell Line and Expression” module in variant detail page of rs4456788. This variant is also considered to repress gene expression in K562 cell line. **C**. The chromatin profile feature plot of rs4456788 focusing on transcription factor binding in HepG2 cell line, sorted by log 2 fold change. The variant is considered to decrease the binding affinity of transcription factors MAZ and FUS.

**Figure S6 Illustration of the protocol details documented in REVA** Protocol details were extracted from original literature and as an attachment in the accession (http://reva.gao-lab.org/download/protocol/reva-0002.pdf).

### Tables

**Table S1 Thresholds of published state-of-the-art tools that we evaluated**

**Table S2 Distribution of locus of positive and negative variants**

**Table S3 Distribution of positive and negative variants on chromosomes**

**Table S4 Annotations of variants with 689 ENCODE profiles**

**Table S5 Information of all ENCODE profiles**

**Table S6 Performance of involved tools on the benchmarking dataset**

**Table S7 Cell line information of benchmark dataset**

**Table S8 Variants in benchmark dataset overlapped with HGMD**

**Table S9 Variants in benchmark dataset overlapped with ClinVar**

### Oligonucleotide library design and synthesis

Oligonucleotide libraries were designed to contain, in order, the universal primer site ACTGGCCGCTTCACTG, the variable 145bp test sequence, Kpnl/Xbal restriction sites (GGTACCTCTAGA), a variable 10-bp tag sequence, and the universal primer site AG ATCGGAAGAGCGTCG. Each sequence was tested with IO unique tags in order to reduce variance due to stochastic rates of amplification of specific plasmids. If a putative enhancer or any of its manipulations contained the recognition sequence for any restriction enzyme (GGTACC, TCTAGA, or GGCCNNNNNGGCC), then that putative enhancer was excluded and an additional one was chosen. The resulting 54,000-plex 200-mer oligonucleotide libraries were synthesized by Agilent, Inc.

### MPRA plasmid construction

Full-length oligonucleotides were isolated using I0% TBE-Urea polyacrylamide gel (lnvitrogen) and then amplified by 20-26 cycles of emulsion PCR using Herculase 11 Fusion DNA Polymerase (Agilent) and primers containing Sfil sites. Purified PCR products were then digested with Sfil (NEB) and directionally cloned into the Sfil-digested MPRA vector pGL4.I0M using One Shot TOPI0 Electrocomp *E. coli* cells (lnvitrogen). To preserve library complexity, the efficiency of transformation was maintained at >3× 10^8^ cfu/mg. The isolated plasmid pool was digested with Kpnl/Xbal to cut between the tested sequence and tag, ligated with a synthetic Kpnl-Xbal fragment containing the SV40 early enhancer/promoter (derived from pGL4.73, Promega) and the *luc2* luciferase ORF (derived from pGL4. 10, Promega) and then transformed into E. coli as described above. Finally, to remove the vector background, the resultant plasmid pool was digested with Kpnl, size selected on a 1% agarose gel, self-ligated and retransformed into *E. coli*.

### Cell culture and transfection

HepG2 cells (ATCC HB-8065) were maintained in Eagle’s Minimum Essential Medium supplemented with I0% fetal bovine serum (FBS) penicillin (50 units/ml) and streptomycin (50 µg/mL). For HepG2 transfections, 5×10^6^ cells were plated in 15-cm **plates. Transfections were performed 24 h after plating using Fugene HD (Promega)** according to the manufacturer’s instructions. In each transfection we used 15 µg of DNA and a Fugene:DNA ratio of 7:2. K562 cells (ATCC CCL-243) were cultured in RPMl-1640 supplemented with 10% FBS and 1% GIBCO Antibiotic-Antimycotic (lnvitrogen). For K562 transfections, 20 mg of DNA was introduced into 4x 10^6^ cells using a Nucleofector II device with Nuclefector Kit Vand program T-016; 24 h post-transfection/nucleofection, cells were lysed in RLT buffer (Qiagen) and frozen at -80^°^C. Total RNA was isolated from cell lysates using RNeasy kit (Qiagen). We chose the transfection method for each cell line that maximized efficiency while minimizing cell death.

### Tag-seq

mRNA was extracted from 100 µg of total RNA using MicroPoly(A)Purist kits (Ambion) and treated with DNase I using the Turbo DNA-free kit (Ambion). First-strand cDNA was synthesized from 400 to 700 ng of mRNA using the High Capacity RNA-to-cDNA kit (Applied Biosystems). Tag-seq sequencing libraries were generated directly from 10% ofa cDNA reaction or 50 ng of plasmid DNA by 26 cycle PCR using Pfu Ultra II HS DNA polymerase 2X master mix (Agilent) and primers. The resultant PCR products were size-selected using 2% agarose E-Gel EX (lnvitrogen). The libraries were sequenced in indexed pools of eight or individually using 36-nt single-end reads on an lllumina HiSeq 2000 instrument.

### Data processing and normalization

To infer the tag copy numbers in each Tag-seq library, all sequence reads were examined, regardless of their quality scores. If the first IO nucleotides of a read perfectly matched one of the 54,000 designed tags, and the remaining nucleotides matched the expected upstream MPRA construct sequence, this was counted as one occurrence of that tag. All reads that did not meet this criterion were discarded. This procedure was repeated separately for the plasmid, HepG2 mRNA, and K562 mRNA pools. The plasmid and mRNA counts for each tag was normalized by the total number of counts from the respective source, and a ratio of the mRNA to plasmid counts was then generated for each tag. A single value was produced for each tested sequence by taking the mean over the tags/replicates, excluding any that bad fewer than 40 plasmid reads. The log_2_ of this value divided by the median was used throughout (this normalization is monotonic and consequently does not affect the P-values for the statistical tests used). Because only a small portion of our tested sequences corresponded to what we later determined to be a functional wild-type enhancer or a nondisruptive mutation, we estimate the 0 baseline level to be approximately the background level of expression for our promoter. Consistent with this, the 2098 sequences with scrambled motifs (and thus no expected expression) have a mean normalized expression of -0.0054 for HepG2 cells and -0.06 for K562 cells. Five probes had 0 RNA counts and their log_2_ values were replaced by - 7 (the smallest non-zero mean had a log_2_ of -6.82).

### Statistical analysis

The paired Wilcoxon signed-rank test is used for comparing different versions of the same set of sequences (e.g., original to scramble). The unpaired Mann-Whitney U-test is used to compare two different sets of sequences (e.g., conserved versus ignoring conservation). Combined P-values are calculated, when indicated, by taking the expression values across multiple factors and using them together for the corresponding statistical test by treating them as one list of values. Where replicates for two sequences are directly compared, we use the individual log replicate values with the unpaired, unequal variance Student’s t-test. Correlations are computed using Pearson’s r, and corresponding permutation P-values are computed as the percentile of the absolute correlation amongst 10 million absolute correlations between the vectors randomly shuffied. P-values are computed in a two-tailed manner, unless otherwise specified.

